# Allosteric Modulation of YAP/TAZ-TEAD Interaction by Palmitoylation and Small Molecule Inhibitors

**DOI:** 10.1101/2023.10.24.563887

**Authors:** Kira R. Mills, Jyoti Misra, Hedieh Torabifard

## Abstract

The Hippo signaling pathway is a highly conserved signaling network that plays a central role in regulating cellular growth, proliferation, and organ size. This pathway consists of a kinase cascade that integrates various upstream signals to control the activation or inactivation of YAP/TAZ proteins. Phosphorylated YAP/TAZ is sequestered in the cytoplasm; however, when the Hippo pathway is deactivated, they translocate into the nucleus, where they associate with TEAD transcription factors. This partnership is instrumental in regulating the transcription of pro-growth and anti-apoptotic genes. Thus, in many cancers, aberrantly hyperactivated YAP/TAZ promotes oncogenesis by contributing to cancer cell proliferation, metastasis, and therapy resistance. Because YAP and TAZ exert their oncogenic effects by binding with TEAD, it is critical to understand this key interaction to develop cancer therapeutics. Previous research has indicated that TEAD undergoes an auto-palmitoylation at a conserved cysteine, and small molecules that inhibit TEAD palmitoylation disrupt effective YAP/TAZ binding. However, how exactly palmitoylation contributes to YAP/TAZ-TEAD interactions and how the TEAD palmitoylation inhibitors disrupt this interaction remains unknown. Utilizing molecular dynamics simulations, our investigation not only provides a detailed atomistic insight into the YAP/TAZ-TEAD dynamics but also unveils that the inhibitor studied influences YAP and TAZ binding to TEAD in distinct manners. This discovery holds significant implications for the design and deployment of future molecular interventions targeting this interaction.

## Introduction

The Hippo signaling pathway is an essential cellular cascade that governs tissue growth and organ size in various organisms, including humans.^1,2^ It plays a pivotal role in the regulation of cell proliferation and apoptosis. In the final step of the Hippo pathway reside Yes-associated protein (YAP) and its paralog, transcriptional co-activator with PDZ-binding motif (TAZ). These proteins serve as key effectors of the pathway, relaying signal transduction to the cell nucleus. The core pathway contains a kinase cascade consisting of the Mammalian Ste20 Kinases (MST1/2) and the adaptor proteins Protein salvador 1 (SAV1) and Mps 1 binder kinase activators (MOB1A/B). Upon activation, the MST1/2-SAV1 complex phosphorylates the Large tumor suppressor kinase (LATS1/2)/MOB1A/B complex. Activated LATS1/2, in turn, phosphorylates YAP/TAZ.^3^ When phosphorylated, YAP/TAZ is sequestered in the cytosol through binding with 14-3-3 proteins and subsequently ubiquitinated and eventually degraded by the proteasome.^4^ In other words, when the Hippo pathway is on, the pathway prevents YAP/TAZ from translocating into the nucleus. This cascade is represented in Figure 1A.

**Figure 1:**
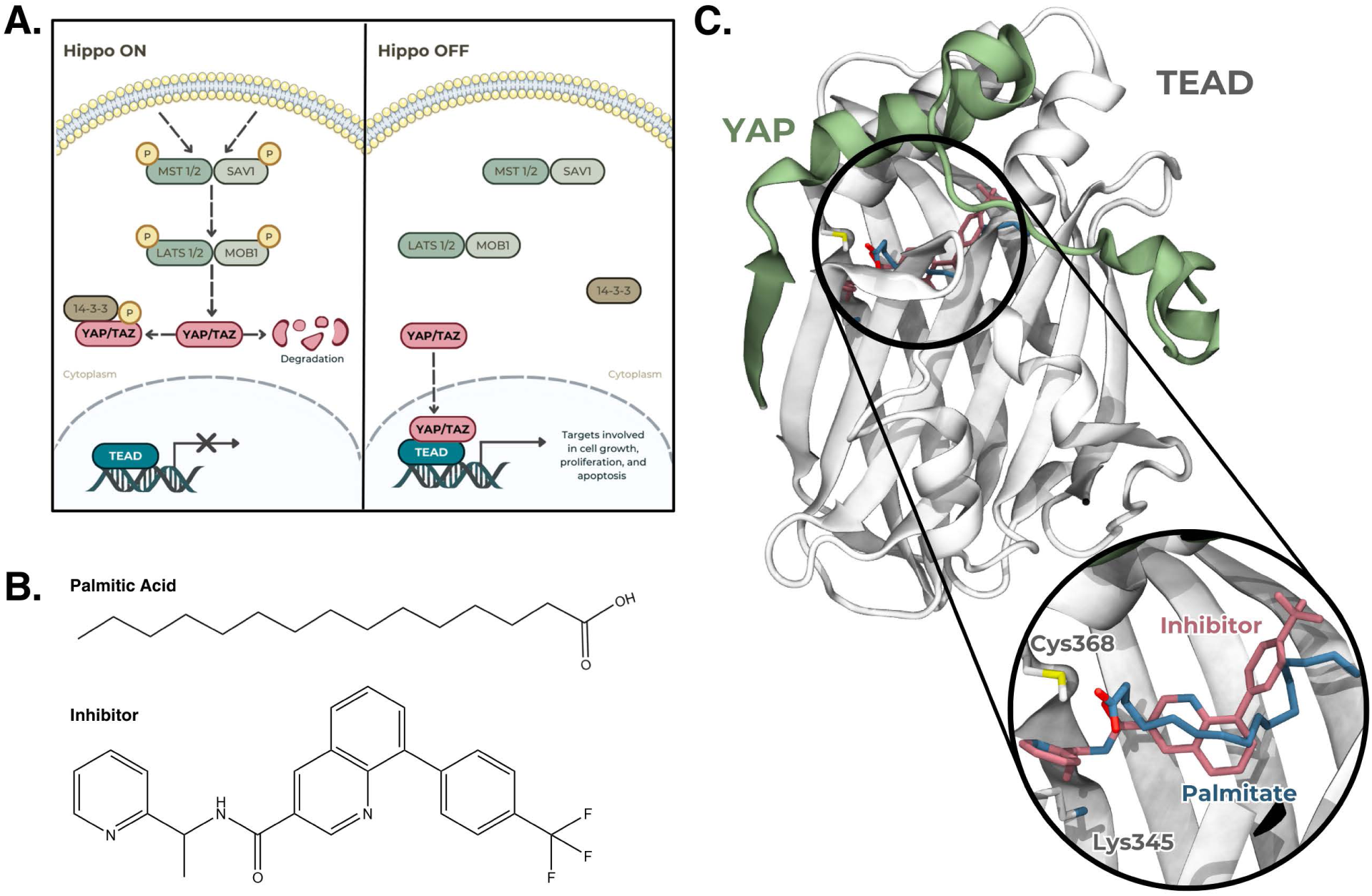
A. A simplified schematic of the Hippo signaling pathway depicting YAP/TAZ in the cytoplasm (when active) and its termination inside the nucleus with YAP/TAZTEAD binding (when turned off). B. Chemical structure of palmitic acid and the inhibitor. C. Crystal structure (PDB ID: 3KYS^10^) of the YAP-TEAD dimer. The palmitate binding pocket is shown in the zoomed-in region, with the conserved Cys368 and Lys345 highlighted, and the known binding positions of palmitate and the inhibitor.

Conversely, when the Hippo pathway is turned off, unphosphorylated YAP/TAZ can translocate into the cell nucleus, where they can bind to the Transcriptional Enhancer Associate Domain (TEAD) family of transcription factors.^5^ Upon binding with TEAD proteins, YAP/TAZ induce the transcription of several target genes linked to cell proliferation and survival (Figure 1A). Consequently, disruption of the Hippo pathway, stemming from mutations in key upstream regulators or dysregulation of its components, leads to the aberrant accumulation and nuclear translocation of YAP and TAZ. Notably, dysregulation of YAP/TAZ-TEAD signaling has been implicated in a spectrum of diseases, including several cancers.^6^

Hence, YAP/TAZ provides an attractive target for therapeutic intervention in cancer treatment. However, YAP and TAZ are not therapeutically amenable, due to their intrinsically disordered structure.^7^ Because YAP and TAZ must associate with TEAD to exert their oncogenic activity, disrupting the YAP/TAZ-TEAD interaction offers an indirect means to inhibit their activity. Additionally, the binding interfaces between YAP/TAZ and TEAD are wide, shallow, and solvent-exposed, making them unsuitable for direct therapeutic targeting.^7^ Consequently, alternative strategies have been devised to allosterically disrupt this interaction.

Because of the crucial role of the Hippo pathway, and specifically of the correctly regulated binding of YAP/TAZ to TEAD, many studies have examined this interaction. ^8–11^ Interestingly, in 2016, Noland *et al* discovered all four TEAD paralogs should be palmitoylated at a conserved cysteine.^12^ Palmitoylation is a known process, in which a palmitate molecule forms a thioester bond with the sidechain of a cysteine residue, impacting signaling and stability.^13^ In the case of TEAD, the palmitate occupies a central hydrophobic palmitate binding pocket (PBP), modulating the stability of TEAD.^12^ In a later study, Chan *et al* demonstrated that palmitate binding and the subsequent palmitoylation of the conserved cysteine had an allosteric effect, enhancing YAP/TAZ association.^14^ The discovery of the PBP and the effect palmitoylation has on TEAD-YAP/TAZ binding, prompted the development of small molecules capable of binding within the PBP, disrupting the YAP/TAZ-TEAD interaction.^15^ One effective inhibitor, VT105, is shown in Figure 1B.

Other studies, however, reported that while TEAD palmitoylation promotes its stability, it was *not* required for YAP/TAZ association.^16,17^ It was further found that a covalent bond might not be necessary between palmitate and cysteine, but instead, palmitate’s presence within the PBP could yield similar effects.^16^ Additionally, recent research suggested that acylation could occur instead at a nearby conserved lysine, resulting in more stabilized TEAD compared to the Cys-palmitoylated protein.^18,19^ One study posits Cys-palmitoylation rendered TEAD in a “transient active form”, while Lys-palmitoylation resulted in a “stable active form”.^19^ Liberelle *et al* recently analyzed available TEAD crystal structures and revealed that the fatty acid could reside freely in the PBP, form a covalent bond with the conserved cysteine, or covalently bind to the conserved lysine.^20^ Despite substantial research on these proteins and their modulation by small molecules, these conflicting results underscore the need for a deeper understanding. Further, the mechanisms by which TEAD palmitoylation and inhibitors allosterically disrupt the YAP/TAZ-TEAD interaction remain elusive.

Here, we have used molecular dynamics (MD) simulations to study the differences in palmitate/palmitic acid and inhibitor binding in the PBP, and how this affects TEAD stability and the YAP/TAZ-TEAD association. Given the conflicting evidence regarding the site of a covalent bond between palmitate and TEAD, our simulations consider all of the small molecules non-covalently bound within the PBP. Additionally, because the pK_a_of palmitic acid^21^ is close to physiological pH (6.37), we ran simulations using both palmitic acid and palmitate. By performing microsecond-long simulations on standalone TEAD, the YAP-TEAD heterodimer, and the TAZ-TEAD heterodimer, in the apo, palmitate-bound, palmitic acid-bound, and inhibitor-bound forms, we produced a total of 36 *µ*s of simulation data. These simulations revealed differences in stability for all three proteins and elucidated distinct impacts of these ligands on YAP and TAZ.

## Computational Details

### System Preparation

The structure of the TEAD3 YAP-binding domain (residues 218-435) was obtained from the RCSB Protein Data Bank (PDB ID: 7CNL^15^). SWISS-MODEL^22^ was used to perform homology modeling and determine appropriate positions for the missing residues 253-260, with a QMEANDisCo global score of 0.83 *±* 0.06 (Figure S1). To determine the protonation state of all titratable residues at a pH of 7.4, the H++ webserver ^23–25^ was used. All Asp and Glu residues were determined to be deprotonated, all Lys residues were determined to be protonated, and all His residues were determined to be protonated at N*ɛ*, except for His250 and His376 which were protonated at N*δ*.

The charges and parameters of the small molecules VT105, palmitate, and palmitic acid were generated using the Antechamber program of AmberTools20 with the AM1-BCC charge scheme and GAFF2.^26^ The crystal structure of the TEAD3 YAP-binding domain used for the protein (PDB ID: 7CNL^15^) also contained VT105 and was thus used for bound VT105 coordinates. To create an apo TEAD monomer, VT105 was removed. To create palmitate and palmitic acid bound systems, the PDB structure 5EMW^12^ was used to align against.

To create the YAP/TAZ-TEAD heterodimer systems, the final snapshots from the TEAD simulations were aligned against either the YAP-TEAD complex structure from PDB ID: 3KYS^10^ or the TAZ-TEAD complex structure from PDB ID: 5GN0^8^ using VMD.^27^

Next, the tleap program of AmberTools20^26^ was used to add missing H atoms. OPC water was added to create a box with a solvent buffer of 15 Å around the protein (approximately 17,200 water molecules), and NaCl ions were added to achieve a salt concentration of 0.15 M, resulting in systems of approximately 70,000 - 85,000 atoms. Once solvated and ionized, tleap was used to generate the initial parameter and coordinate files for each of the systems, using Amber’s ff19SB,^28^ and OPC^29^ force fields.

### Molecular Dynamics Simulations

All simulations were conducted using the pmemd.cuda implementation of the Amber20 software package.^26,30,31^

To ensure proper relaxation and minimization of the structures, a multistep minimization procedure was used, each of which comprised 5000 steps of steepest descent minimization. First, only the water and ion atoms were relaxed, with a 5 kcal/mol/Å^2^ restraint placed on all other atoms. Second, the protein sidechains were also allowed to relax, with a 5 kcal/mol/Å^2^ restraint placed on only the protein backbone. Lastly, the entire system was allowed to relax, by removing all restraints.

After minimization, the systems were heated to 300 K over 1 ns using the Langevin thermostat,^32^ during which time the protein backbone was held with a 1 kcal/mol/Å^2^ restraint. Next, an NPT equilibration step was performed at 300 K with a 1 kcal/mol/Å^2^ restraint placed on the protein backbone. For the TEAD-only systems, this equilibration was run for 10 ns, and for the larger dimer systems, it was extended to 20 ns.

Finally, 1 *µ*s of production simulations were run for each of the systems, repeated in triplicate. A time step of 2 fs was used, and all bonds involving hydrogen were constrained using the SHAKE algorithm.^33^ The cutoff distance for long-range interactions was set to 12 Å. The simulations were performed in an isothermal and isobaric ensemble, with the temperature held at 303 K, using the Langevin thermostat^32^ and Berendsen barostat.^34^

### Analysis

All simulation analyses except EDA were performed using the cpptraj program from the Amber20 software package.^35^ Root mean square deviation (RMSD) was calculated for the backbone atoms. Root mean square fluctuation (RMSF) was calculated for non-hydrogen sidechain atoms. The interaction energies were calculated and reported as the sum of the electrostatic and van der Waals energies using the linear interaction energy (LIE) function in cpptraj.

To determine protein-ligand contacts, cpptraj was again used to find the simulation fractions during which residues were in contact with the ligand, with a contact being defined as having atoms within 7 Å of each other. To identify specific modes of interaction between the protein and ligand, the Protein Ligand Interaction Profiler (PLIP) web tool was used.^36^ For each simulation, snapshots were taken every 100 ns and uploaded to the PLIP web tool, which then identified any hydrophobic interactions, hydrogen bonds, salt bridges, *π*-stacking, *π*-cation interactions, water bridges, halogen bonds, or metal complexes.

To perform Energy Decomposition Analysis (EDA) a Fortran code available on GitHub was used.^37^ EDA calculates the average pairwise intermolecular van der Waals and Coulomb interactions as a function of a residue of interest. This analysis allows us to assess the roles of specific amino acid residues in stabilizing or destabilizing the protein-protein interactions between TEAD and YAP or TAZ. This analysis has been employed several times previously for QM/MM and MD simulations of protein systems.^38–41^

## Results and Discussion

### Effect of PBP ligands on the dynamics of TEAD

Initially, all-atom MD simulations were run on four systems containing the TEAD YAP-binding domain: the favored palmitic acid- and palmitate-bound systems, and the unfavored apo and inhibitor-bound systems. All of the small molecules bind in the PBP, containing the conserved Cys (Cys368) and Lys (Lys345) which undergo the auto-palmitoylation reaction, as shown in Figure 1C.

For all of these initial TEAD systems, 1 *µ*s simulations were run in triplicate, and the combined results are displayed in Figure 2. In Figure 2A, the RMSD of the TEAD backbone during each of the simulations is presented, highlighting that upon small molecule binding, the inhibitor and palmitic acid exhibit similar levels of TEAD stabilization as evidenced by the decreased RMSD. Compared to the apo form, palmitate-bound TEAD shows slight stabilization, albeit not as pronounced as the other two small molecules, and with a more bimodal distribution. This suggests, overall, that TEAD is stabilized when small molecules bind in the PBP, which could be beneficial for the subsequent binding of YAP or TAZ.

**Figure 2:**
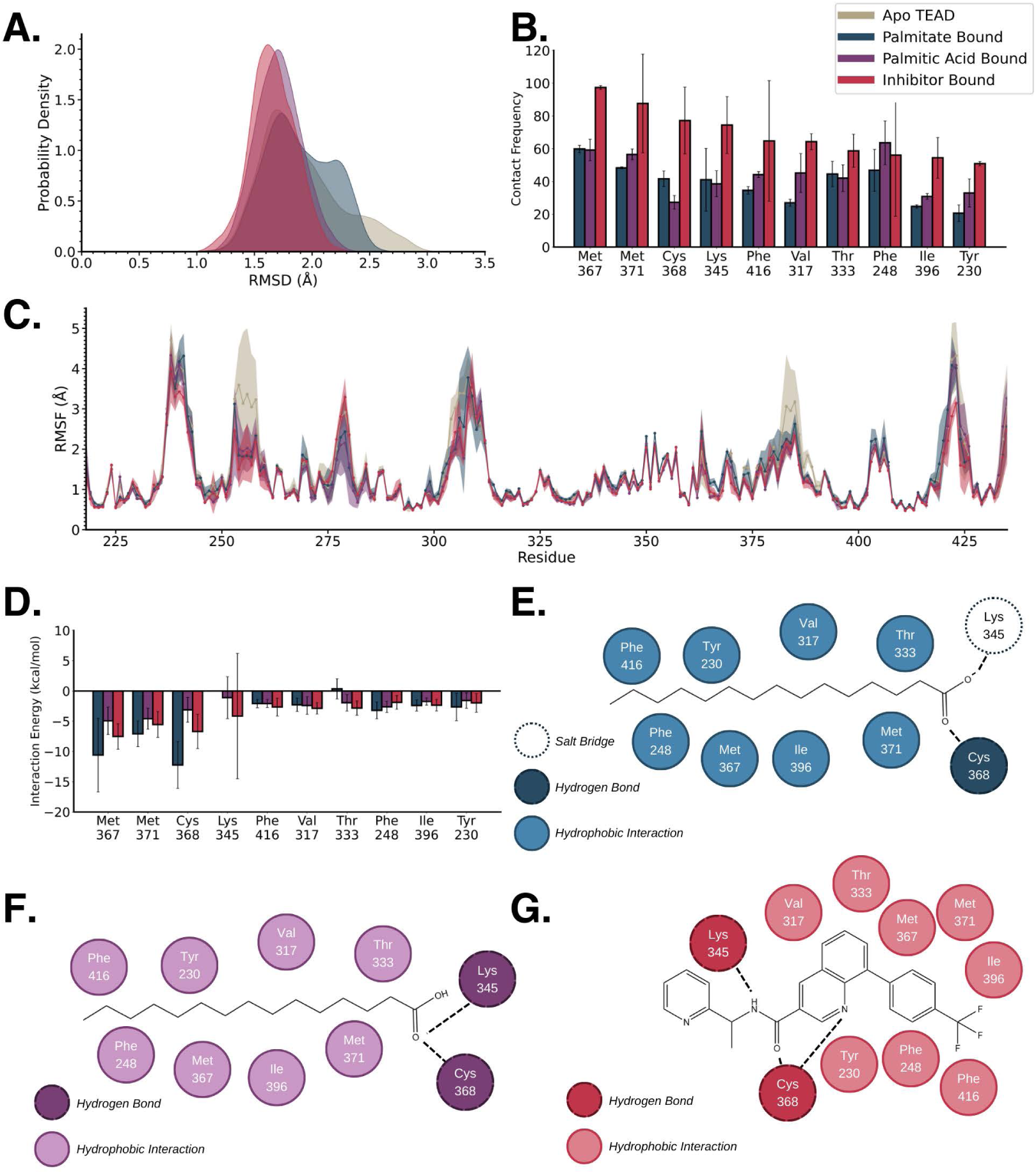
Combined results for simulations of TEAD only. A. RMSD of TEAD backbone atoms in each of the four systems studied. B. Average contact frequency between each of the small molecules and the residues lining the palmitate binding pocket. Error bars show the standard deviation between the three trials. C. RMSF of TEAD residues during the simulations. Solid lines show average values and transparent areas show standard deviation. D. Average interaction energy between each of the small molecules and the residues lining the palmitate binding pocket. Error bars show the standard deviation between the three trials. The interaction energy between palmitate and Lys345 is omitted here to better visualize other results but is shown in the SI. E-G. Interaction maps classifying the type of interaction between the residues lining the binding pocket and E. palmitate, F. palmitic acid, or G. inhibitor.

The contact frequency between the small molecules and the residues lining the binding pocket, shown in Figure 2B, indicates that the inhibitor consistently maintains more contacts compared to either palmitate or palmitic acid. It is also the only one that shows *>*90% contacts with any residues throughout the simulation, including the two possible sites of palmitoylation, Cys368 and Lys345. This suggests that the inhibitor binds more tightly, with less movement and fluctuation inside the binding pocket. This observation is further explored in subsequent panels.

To delve deeper into these differences in protein motion, the RMSF of the TEAD residue sidechains is shown in Figure 2C. In this figure, several regions stand out with large differences between the four systems. Specifically, two regions show increased fluctuation in the apo TEAD system compared to the other three: residues 253-258 and 382-386. Similarly, two regions highlight the inhibitor-bound TEAD as having significantly less fluctuation than the other three: residues 238-241 and 420-426. Although none of these regions comprise the palmitate-binding pocket, two of them (382-386 and 420-426) are part of the YAP/TAZ binding interfaces which will be explored further in subsequent sections. Because these regions necessary for successful protein-protein interactions exhibit increased fluctuation in the apo TEAD and decreased fluctuation in the inhibitor-bound TEAD, these results indicate that some degree of stabilization from small molecule binding is favorable for subsequent protein binding. However, the inhibitor is capable of over-stabilizing TEAD, making YAP/TAZ association unfavorable.

The average interaction energy between the small molecules and the residues lining the binding pocket is presented in Figure 2D. Here, palmitate consistently exhibits stronger interaction energies, despite having less frequent contacts, as shown in Figure 2B. This can be attributed to the fact that the interaction energies shown are the sum of van der Waals and electrostatic energies. Consequently, the negatively charged palmitate can form stronger interactions. As a note, the interaction energy between palmitate and Lys345 is omitted in Figure 2D but is shown in Figure S2. This omission is due to the large interaction energy between Lys345 and palmitate, which overshadowed the other data when included in the same figure. When comparing only the inhibitor and palmitic acid, the inhibitor exhibits stronger interactions, including with Cys368 and Lys345, consistent with the results presented in Figure 2B as the inhibitor can bind more tightly.

In Figures 2E-G, the interaction maps clarify the type of interaction each small molecule forms with the residues lining the binding pocket. These maps reveal similarities, with most residues forming hydrophobic interactions with the small molecules. However, palmitate is capable of forming a salt bridge with Lys345 (Figure 2E), explaining the large interaction energy shown in Figure S2. The primary distinction between the inhibitor and the native ligands lies in where they form stronger interactions, such as hydrogen bonds or salt bridges. Due to the location of their polar groups, both palmitate and palmitic acid can only form these interactions at the terminal carboxylate/carboxylic acid (Figures 2E and F), where they interact with the sidechains of Cys368 and Lys345. In contrast, the inhibitor can form hydrogen bonds with those same residues on more central atoms (Figure 2G). This could contribute to the inhibitor’s more consistent retention within the pocket. Additionally, the inhibitor’s larger and more three-dimensional structure provides better opportunities for contact with the residues lining the pocket.

Previous studies have demonstrated that the binding and reaction of palmitate/palmitic acid enhance TEAD’s ability to bind YAP or TAZ. These initial simulation results indicate a need for some flexibility in TEAD for subsequent protein binding. When the inhibitor binds as tightly as it does, it reduces TEAD’s overall flexibility, which may contribute to its inhibitory capability. However, in the absence of any small molecules, TEAD may be too flexible, contributing to why the apo TEAD has reduced success in YAP/TAZ binding. Furthermore, because the inhibitor does have better interactions with the residues of TEAD, this may explain its ability to out-compete the native substrates.

Following these simulations studying TEAD alone with small molecules, we aimed to understand how this could affect the subsequent binding of YAP or TAZ. To achieve this, we obtained final snapshots from the TEAD simulations and bound either YAP or TAZ before performing another 1 *µ*s of simulations in triplicate.

### Effect of PBP ligands on the dynamics of YAP-TEAD heterodimer

The combined results for the systems with YAP bound are summarized in Figure 3. The RMSDs of the backbone atoms of both TEAD and YAP proteins are presented in Figures 3A and 3B, respectively. In both, there is a noticeable reduction in protein movement in the unfavored apo and inhibitor-bound systems compared to the favored palmitate- or palmitic acid-bound systems. This difference is more pronounced for YAP but is still evident for TEAD as well.

**Figure 3:**
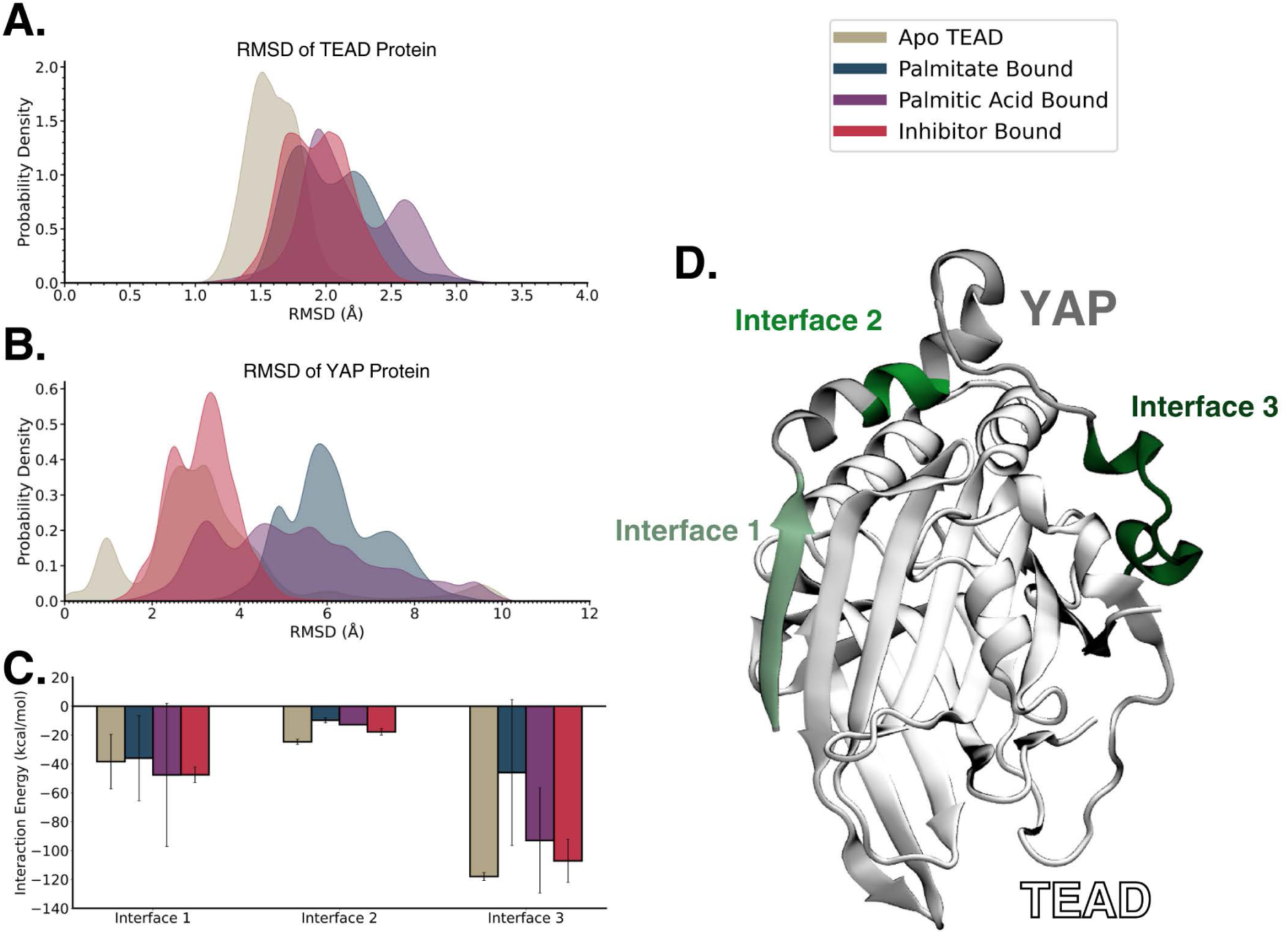
Combined results for the YAP-TEAD heterodimer simulations. A. RMSD of the backbone atoms of TEAD protein in each of the simulations. B. RMSD of the backbone atoms of YAP protein in each of the simulations. C. Total interaction energy between the residues comprising each of the binding interfaces of YAP and TEAD. Error bars show the standard deviation between the three trials. D. Visualization of the three binding interfaces between YAP and TEAD.

Focusing on the TEAD RMSD (Figure 3A), in the TEAD-only systems (Figure 2A) the RMSD for the apo system exhibited a broader distribution with peaks near 1.8 Å and 2.5 Å. After YAP binding, there is a narrowing of this distribution, with a peak instead occurring at 1.5 Å. This shift indicates TEAD stabilization upon YAP binding, although in this unfavorable system, it could potentially be an overstabilization. In the inhibitor-bound system, there is instead a slight broadening of the RMSD distribution from the TEAD-only system. However, the peak remains centered around 1.8 Å, indicating less pronounced changes in TEAD behavior. The palmitate-bound system similarly shows little change upon YAP binding. In ctonrast, the palmitic acid-bound system exhibits a significant broadening of its RMSD distribution and a shift to a bimodal distribution upon YAP binding.

Examining the YAP RMSD (Figure 3B), significant differences are observed between the favored (palmitate- and palmitic acid-bound) and unfavored (apo and inhibitor-bound) systems. The former show RMSD distributions that are much broader and have peaks at higher RMSDs compared to the latter. The apo and inhibitor-bound systems show peaks around 3 Å, whereas the palmitate- and palmitic acid-bound systems exhibit peaks between 3 Å and 8 Å, respectively. This indicates that increased flexibility in YAP likely improves its ability to interact correctly with TEAD.

To examine the areas where YAP and TEAD have been previously reported to interact, Figure 3C presents the total interaction energies between the two proteins, focusing on the residues involved in one of the three binding interfaces, ^10^ which are visualized in Figure 3D. Examining all interaction energies, there are no significant differences between the systems within each interface, with the exception of the palmitate-bound system at Interface 3, which exhibits weaker interaction with TEAD compared to the other three systems. Generally, the apo and inhibitor-bound systems have slightly stronger interaction energies than the palmitate- or palmitic acid-bound systems. This likely contributes to the decreased movement of both YAP and TEAD observed in the apo and inhibitor-bound systems, as discussed with their backbone RMSDs.

In Figure 4, the EDA is displayed for each YAP residue with the entire TEAD protein. The values represent the combined van der Waals and electrostatics energies and are displayed as the difference between the systems with palmitate, palmitic acid, or the inhibitor bound and the apo system, illustrating whether the binding of the small molecules stabilizes or destabilizes the YAP-TEAD interactions. These Δ*E* values are also presented in Table S1, and the original, unsubtracted values are shown in Figure S3. Overall, the EDA results align with the trends seen in the interface interaction energies in Figure 3C.

**Figure 4:**
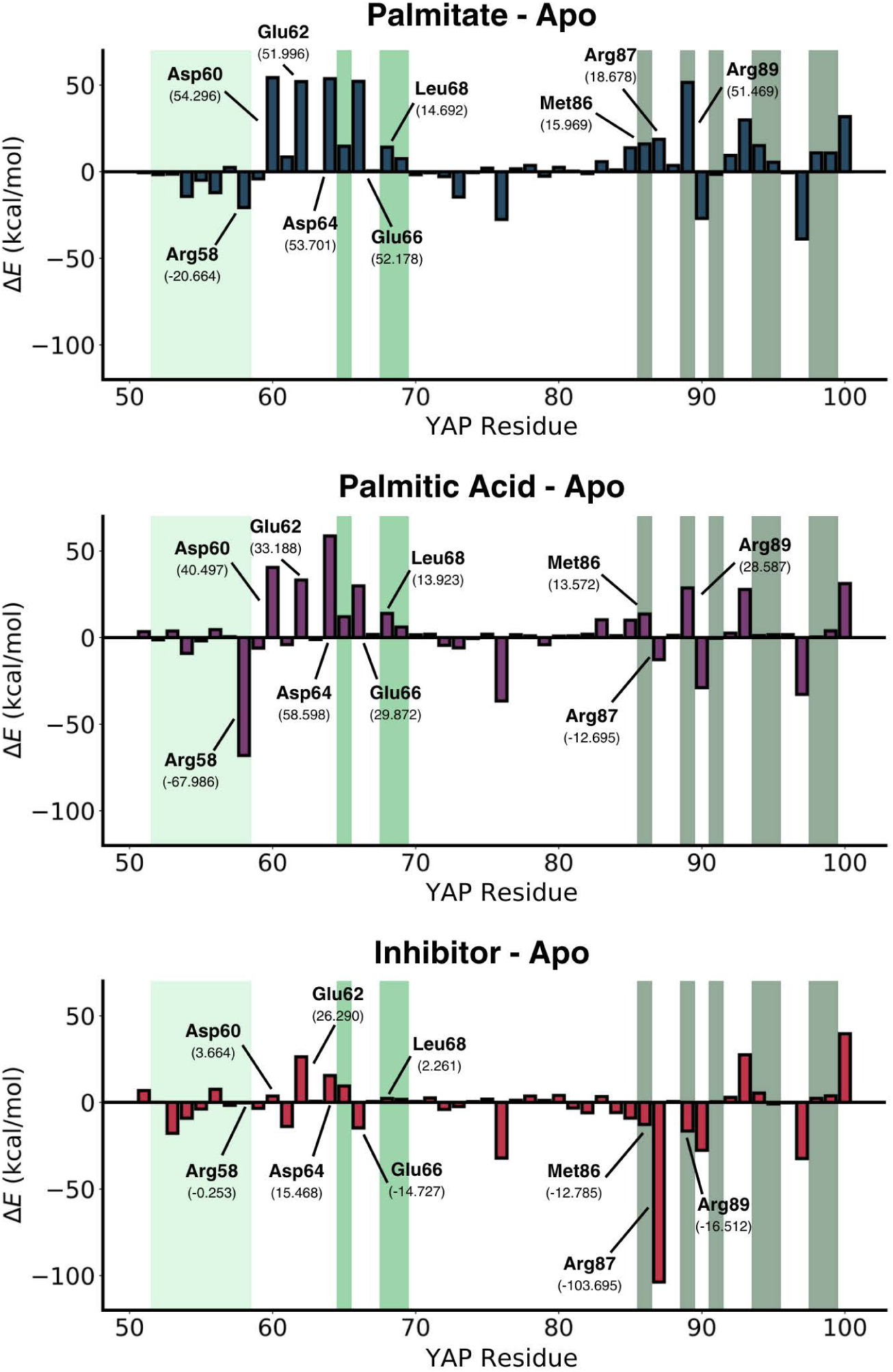
EDA analysis between individual YAP residues and the combined TEAD protein. For each residue, the average delta energy is shown between the systems with palmitate, palmitic acid, or the inhibitor bound and the apo system to show if the binding of the small molecule is stabilizing or destabilizing the interaction between YAP and TEAD. Error bars show the standard deviation between the trials. Values are also given in Table S1. Distinct shaded regions highlight the three binding interfaces. The lightest shade of green marks Interface 1 (residues 52-58), an intermediate shade of green indicates Interface 2 (residues 65, 68, 69), and the darkest shade represents Interface 3 (residues 86, 89, 91, 94-95, 98-99).

Within the YAP residues of Interface 1 (residues 52-58), slight differences are noted between the systems’ interactions in residues 52-57. However, the most significant difference is in residue Arg58. In the unfavored inhibitor-bound system, there is almost no change from the apo system’s original repulsion, whereas the favored systems both experienced stabilization (Δ*E* = *−*20.664 and Δ*E* = *−*67.986 kcal/mol for palmitate- and palmitic acid-bound, respectively). In Interface 2 (residues 65, 68, 69), all systems exhibit a slight destabilization at Leu68, with the inhibitor-bound system showing the smallest change. Finally, in Interface 3 (residues 86, 89, 91, 94-95, 98-99), the primary difference is observed in residues Met86 and Arg89. In both palmitate- and palmitic acid-bound systems, these residues demonstrate destabilization (combined Δ*E* = 67.438 and Δ*E* = 42.159 kcal/mol, respectively), while the inhibitor-bound system instead showed stabilization (combined Δ*E* = *−*29.297 kcal/mol).

Moreover, focusing on residues outside the binding interfaces, a noticeable series of acidic residues (Asp60, Glu62, Asp64, and Glu66) exhibit varying behavior between the three systems. In the palmitate-bound system, all four residues almost equally destabilize the YAP-TEAD interaction, with Δ*E* = 52 *−* 54 kcal/mol for each. In the palmitic acid-bound system, all four residues still contribute to destabilization, although the Asp residues have a more significant impact than the Glu residues. In comparison to the two favored systems, the inhibitor-bound system shows a much less pronounced destabilizing effect. Except Glu66 which is stabilized (Δ*E* = *−*14.727 kcal/mol) in this system. Additionally, Arg87 shows moderate destabilization in the palmitate-bound system (Δ*E* = 18.678 kcal/mol), and moderate stabilization in the palmitic acid-bound system (Δ*E* = *−*12.695 kcal/mol). However, in the inhibitor-bound system, it exhibits significant stabilzation (Δ*E* = *−*103.695 kcal/mol). Finally, when considering all YAP residues together by summing the Δ*E* values, the unfavored inhibitor-bound system experiences significant stabilization (Δ*E* = *−*152.956 kcal/mol), while the favored palmitate- and palmitic acid-bound systems instead undergo substantial destabilization (Δ*E* = 315.274 and Δ*E* = 134.485 kcal/mol, respectively), consistent with the RMSD data.

These EDA results support the work by Li *et al*, which found that mutations of Met86 and Arg89 led to significant reductions in the ability of YAP and TEAD to associate, and mutation of Leu68 resulted in reduced function of the YAP-TEAD system.^10^ Outside of the binding interfaces, overall, there are stronger interaction energies, mainly attractive, between apo and inhibitor-bound TEAD with YAP, which could be a contributing factor to the reduced movement of both proteins observed in those systems’ RMSD.

### Effect of PBP ligands on the dynamics of the TAZ-TEAD heterodimer

Similarly, the combined results for the systems with TAZ bound are summarized in Figure 5. One simulation has been excluded from these results, which is a palmitic acid-bound TAZ-TEAD heterodimer trial, because the palmitic acid left the PBP, subsequently causing TAZ to dissociate. However, snapshots and analyses from this trial are included in Figures S4 and S5.

**Figure 5:**
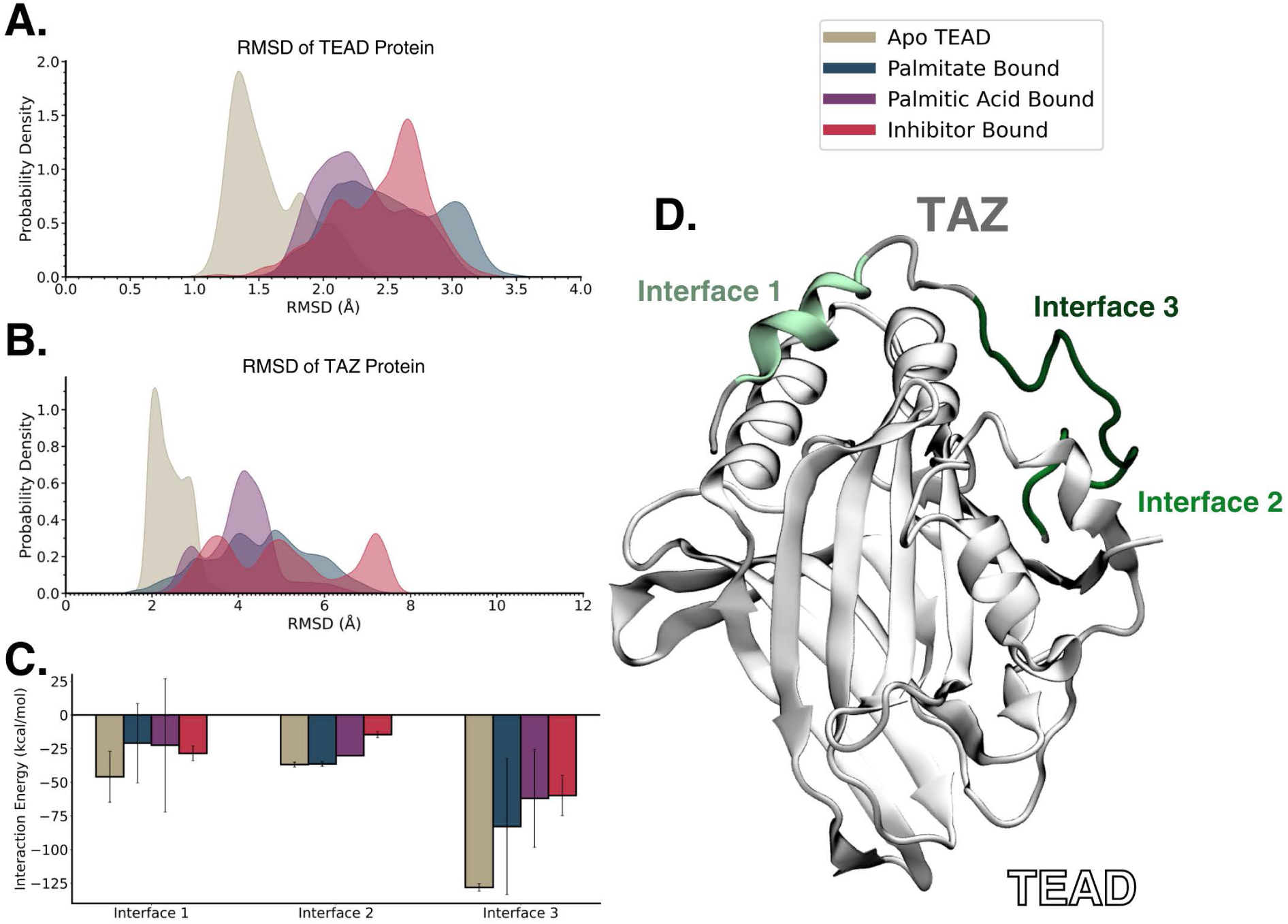
Combined results for the TAZ-TEAD heterodimer simulations. One palmitic acidbound trial is omitted, data including this trial is shown in Figure S5. A. RMSD of the backbone atoms of TEAD protein in each of the simulations. B. RMSD of the backbone atoms of TAZ protein in each of the simulations. C. Total interaction energy between the residues comprising each of the binding interfaces of TAZ and TEAD. Error bars show the standard deviation between the trials. D. Visualization of the three binding interfaces between TAZ and TEAD.

In Figure 5A, the RMSD of the backbone atoms of TEAD are shown. Much like what occurred for apo TEAD upon binding of YAP, once bound to TAZ there is a narrowing of the RMSD distribution, and a slight decrease at the peak location, from 1.8 Å and 2.2 Å to 1.4 Å. Again, since this is an unfavorable system, and its RMSD is clearly lower than the other systems, it appears that the stabilization resulting from TAZ binding may potentially represent an overstabilization of TEAD. However, when comparing the other three systems, there is a substantial overlap between them with less clear differentiation. The palmitic acid-bound system exhibits a slightly narrower distribution and a lower peak compared to either the palmitate- or inhibitor-bound system. Nevertheless, when compared to their respective TEAD-only behaviors, both the inhibitor- and palmitate-bound systems demonstrate noticeable broadening of their distributions and increases in the peak locations, indicating increased TEAD fluctuations after TAZ binding. For the palmitic acid-bound system, there is much less broadening but still a slight increase in the peak location.

Examining the RMSD of the TAZ backbone in Figure 5B, the systems display a similar trend but with more apparent differences. Once again, the apo system exhibits the lowest RMSD, indicating increased stabilization compared to the other three systems. For the inhibitor-bound system, unlike the YAP bound system, when TEAD binds, the TAZ protein shows the highest RMSD and broadest distribution, indicating significantly more fluctuation than any of the other systems. This could suggest that, while the inhibitor can disrupt the YAP-TEAD dimer’s function by overstabilization, it may disrupt the TAZ-TEAD interaction by preventing proper protein-protein interactions.

To further investigate these protein-protein interactions, Figure 5C presents the total interaction energies between the two proteins, focusing on the residues involved in one of the three binding interfaces, which are visualized in Figure 5D. It’s worth noting that while the three binding interaces were previously described for YAP-TEAD,^10^ only two interfaces have been described for TAZ-TEAD, and what we refer to as Interface 3 represents a series of crucial hydrogen bonds described by Kaan *et al*.^8^ These interaction energies clearly demonstrate a decreasing trend in interaction strength in Interfaces 2 and 3, with apo *>* palmitate-bound *>* palmitic acid-bound *>* inhibitor-bound systems. This trend aligns with the earlier observation that the apo system might be binding TAZ too tightly, resulting in decreased movement. In contrast, the inhibitor-bound system may not bind TAZ tight enough.

In Figure 6 the results of the EDA are presented for each residue of TAZ with all TEAD residues. These values represent the sum of the van der Waals and electrostatic interactions and are displayed as the difference between each of the small molecule bound systems and the apo system, revealing how the small molecules stabilize or destabilize the TAZ-TEAD interactions. These Δ*E* values are also shown in Table S2, and the original, unsubtracted values are shown in Figure S6.

**Figure 6:**
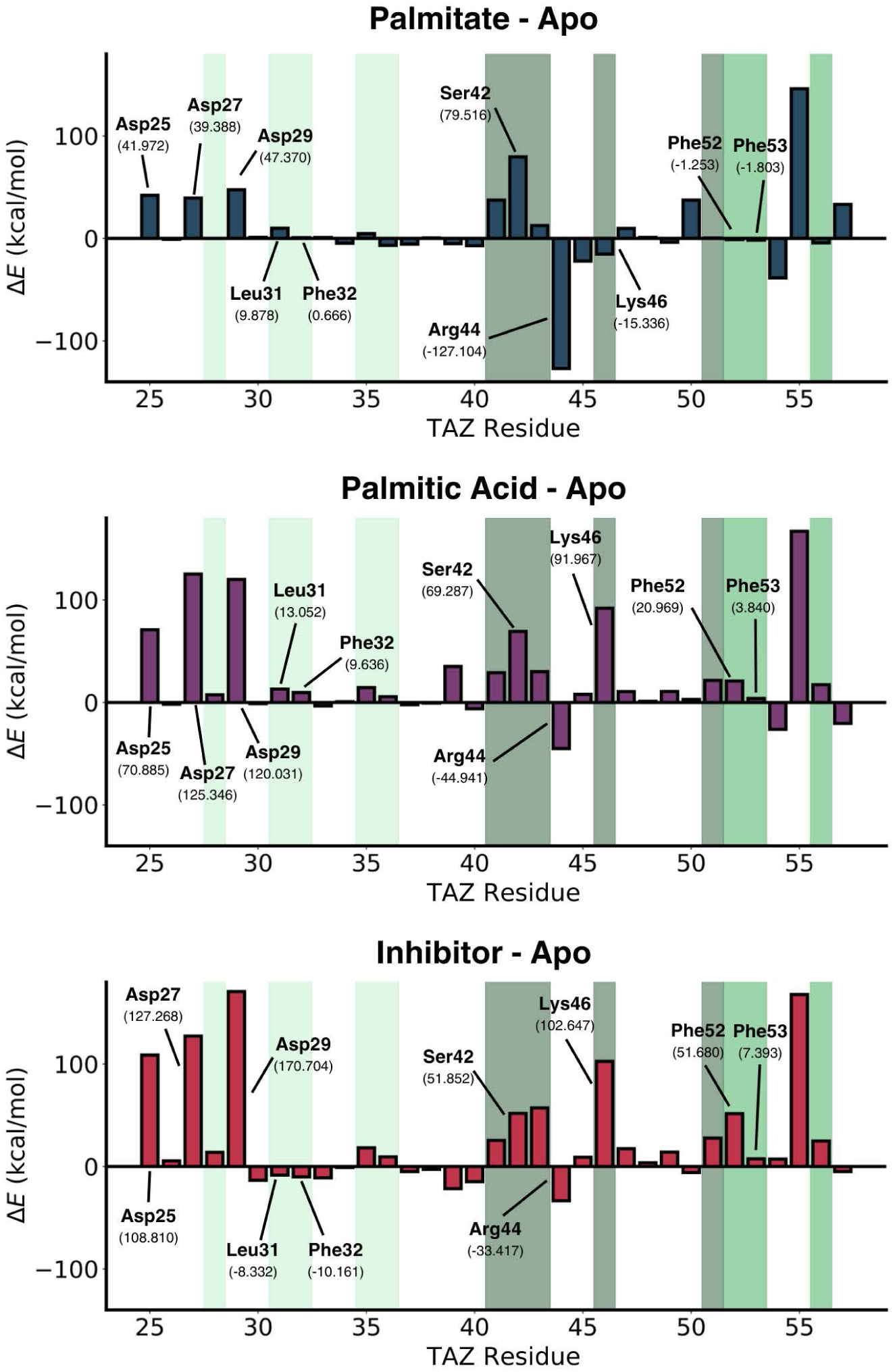
EDA analysis between individual TAZ residues and the combined TEAD protein. For each residue, the average delta energy is shown between the systems with palmitate, palmitic acid, or the inhibitor bound and the apo system to show if the binding of the small molecule is stabilizing or destabilizing the interaction between TAZ and TEAD. Error bars show the standard deviation between the trials. Values are also given in Table S2. Distinct shaded regions highlight the three binding interfaces. The lightest shade of green marks Interface 1 (residues 28, 31-32, 35-36), an intermediate shade of green indicates Interface 2 (residues 52, 53, 56), and the darkest shade represents Interface 3 (residues 41-43, 46, 51). Palmitic acid-bound trial that showed dissociation is omitted as EDA analysis gave infinite values.

Similar to the YAP-TEAD systems, the overall trends seen in the EDA align with the trends shown in the interaction energies of Figure 5C for the binding interfaces. Within Interface 1, there is little difference between the systems, and all of them show relatively small changes from the apo system. However, one area that stands out is Leu31 and Phe32, where both palmitate- and palmitic acid-bound systems exhibit moderate destabilization, while the inhibitor-bound system shows slight stabilization. In Interface 2, residues Phe52 and Phe53 display a clear increase in destabilization from palmitate-bound (Δ*E* = *−*1.253 and Δ*E* = *−*1.803 kcal/mol) to palmitic acid-bound (Δ*E* = 20.969 and Δ*E* = 3.840 kcal/mol) to inhibitor-bound (Δ*E* = 51.680 and Δ*E* = 7.3934 kcal/mol). Finally, in Interface 3, there are more pronounced differences between the systems. Looking first at Ser42, both the favored palmitate- and palmitic acid-bound systems exhibit more substantial destabilization (Δ*E* = 79.516 and Δ*E* = 69.287 kcal/mol, respectively) compared to the inhibitor-bound system (Δ*E* = 51.852 kcal/mol). Interestingly, in Lys46, the favored palmitic acid-bound system (Δ*E* = 91.967 kcal/mol) behaves more similarly to the unfavored inhibitor-bound system (Δ*E* = 102.647 kcal/mol), than it does to the palmitate-bound system, which undergoes stabilization (Δ*E* = *−*15.336 kcal/mol).

Furthermore, when looking outside of the binding interfaces, more differences emerge between the systems. First, much like what was observed with the YAP-TEAD results in Figure 4, there is a series of early acidic residues in TAZ (Asp25, Asp27, and Asp29) that show different levels of destablization after small molecule binding. Unlike with the YAP acidic residues, however, the acidic residues in TAZ show higher destabilization in the inhibitor-bound system (total Δ*E* = 406.782 kcal/mol) than in either the palmitate- or palmitic acid-bound systems (Δ*E* = 128.730 and Δ*E* = 316.262 kcal/mol, respectively). Additionally, Arg44 undergoes higher stabilization in the favored systems (palmitate-bound Δ*E* = *−*127.104 and palmitic acid-bound Δ*E* = *−*44.941 kcal/mol) than in the unfavored inhibitor-bound system (Δ*E* = *−*33.417 kcal/mol). Finally, when all of the TAZ residues are considered together, these results demonstrate the same trend observed earlier, where the amount of destabilization for each of the small molecules is palmitate-bound (Δ*E* = 258.132 kcal/mol) *<* palmitic acid-bound (Δ*E* = 779.231 kcal/mol) *<* inhibitor-bound (Δ*E* = 889.467 kcal/mol).

These results are consistent with those shown in Figure 5, providing evidence for the inhibitor-bound system’s apparent inability to bind TAZ as tightly as the other systems can. Among the highlighted residues, mutations of several have been experimentally shown to affect proper protein-protein interactions. Chan *et al* ^42^ demonstrated that paired mutations of Asp27-Leu28, Leu31-Phe32, Trp43-Arg44, and Phe52-Phe53 reduced the ability for TAZ-TEAD association. Additionally, Zhang *et al* ^43^ highlighted the importance of Ser51, which did not stand out in our results, but which did follow the same trend observed above.

## Conclusions

In this study, we employed molecular dynamics (MD) simulations to investigate the impact of small molecule binding on TEAD proteins in isolation and in the context of YAP-TEAD and TAZ-TEAD heterodimers. Previous experimental studies have proposed an auto-palmitoylation reaction within the palmitate-binding pocket (PBP) as a crucial step for successful YAP/TAZ-TEAD binding. However, this conclusion has been debated in the literature.^12,14,16–20^ Following the discovery of the PBP, several groups have developed small molecule inhibitors designed to occupy this region and prevent palmitoylation. ^15,44–48^ In this work, we examined the dynamics of apo TEAD, or TEAD bound to either palmitate/palmitic acid, or the inhibitor VT105. Our simulations revealed that the binding of any of these small molecules has a stabilizing effect on TEAD, and the inhibitor had a higher propensity to remain in the PBP compared to the other small molecules.

From our YAP-TEAD simulations, it became evident that the unfavored apo and inhibitor-bound systems exhibited increased stabilization of both TEAD and YAP, compared to the palmitate- or palmitic acid-bound systems. This observation suggests that a certain degree of conformational flexibility in the proteins is necessary for the formation of the correct and functional YAP-TEAD heterodimer. Moreover, we identified key differences in how YAP and TEAD associate with specific YAP residues, including Arg58, Asp60, Glu62, Asp64, Glu66, Leu68, Met86, Arg87, and Arg89. Previous mutagenesis studies on YAP have reported that mutations of Met86 and Arg89 led to significant decreases in the ability of YAP and TEAD to associate.^10^ Additionally, mutations of Leu68 resulted in reduced function of the YAP-TEAD system.^10^

Conversely, our TAZ-TEAD simulations showed similar stabilization of the apo systems but increased flexibility in the inhibitor-bound system. We highlighted the TAZ residues Leu31, Phe32, Ser42, Arg 44, Lys46, Phe52, and Phe53, as being significantly affected by the presence of small molecules in their ability to associate with TEAD. A previous study investigated the double mutations of Asp27-Leu28, Leu31-Phe32, Trp43-Arg44, and Phe52-Phe53,^42^ which reduced the ability for TAZ-TEAD association. However, the other residues we emphasized have not been studied experimentally, suggesting potential targets for future research to determine their roles in the TAZ-TEAD interaction.

Collectively, our findings indicate the existence of a flexibility sweet spot in which TEAD and YAP/TAZ must exist for proper function. The inhibitor VT105 appears to act on them differently, potentially keeping them out of this optimal state and disrupting their interaction. This study provides a mechanistic insight into how the PBP ligands allosterically modulate the YAP/TAZ-TEAD interactions, with direct implications for the successful computational prediction of small molecule inhibitors that bind the PBP and disrupt these interactions. Notably, molecules that simply bind to the PBP may not necessarily disrupt YAP/TAZ-TEAD interactions. Therefore, successful computational predictions should consider how small molecules affect the critical residues identified in this study as key modulators of YAP/TAZ-TEAD interactions. Furthermore, these results can guide future hit-to-lead optimization during drug development by predicting how different analogs affect these residues. Intriguingly, our study revealed that the inhibitor VT105 affects YAP and TAZ differently. This differential response to the same small molecule adds to the growing body of evidence that, despite their many similarities and previous treatment as redundant, YAP and TAZ exhibit differences in expression, activation, and function.^49–53^ Moreover, crystal structures of YAP-TEAD and TAZ-TEAD complexes have revealed that TAZ-TEAD can engage in an additional heterotetramer binding mode not observed with YAP-TEAD.^8^ In this unique binding mode, the previously identified binding Interfaces 1 and 2 remain unchanged, but Interface 3 loses all of its hydrogen bonds except that of Trp43.^8^ While our study focused on the TAZ-TEAD heterodimer binding mode, which closely resembles the known YAP-TEAD binding mode, future investigations concentrating on the heterotetramer will provide valuable additional insights.

## Data and Software Availability

Simulation files, including parameter, initial coordinate, and trajectory files are available at https://doi.org/10.5281/zenodo.10020092. The Amber20 software used to run molecular dynamics simulations and AmberTools20 software used to build Amber topologies and perform analysis are available on their website (ambermd.org). The PLIP web tool used to identify protein-ligand interactions, EDA software for calculating protein-protein interaction energies, and VMD used for visualization are also freely available at their respective websites (https://plip-tool.biotec.tu-dresden.de/plip-web/plip/index, https://doi.org/10.5281/zenodo.4469902, www.ks.uiuc.edu/Research/vmd/).

## Author Contributions

H.T. and J.M. designed the project; H.T. supervised the project; K.M. performed the computational studies and analyses; H.T. and K.M. analyzed the data; H.T., J.M. and K.M. wrote the manuscript together.

## Acknowledgement

The authors acknowledge the Office of Information Technology Cyberinfrastructure Research Computing (CIRC) at The University of Texas at Dallas for providing HPC resources that have contributed to the research results reported within this paper. This work was funded by startup funds from The University of Texas at Dallas.

## Supporting Information Available

Supporting Information: Tables showing the Δ*E* EDA values for YAP-TEAD and TAZ-TEAD heterodimer systems; results of TEAD homology modeling used to add missing residues; average interaction energy between small molecules and TEAD residues lining the PBP; original, unsubtracted EDA values for YAP-TEAD and TAZ-TEAD heterodimer systems; simulation snapshots showing TAZ dissociation in omitted TAZ-TEAD heterodimer trial; combined TAZ-TEAD results including the omitted trial (PDF)

## TOC Graphic

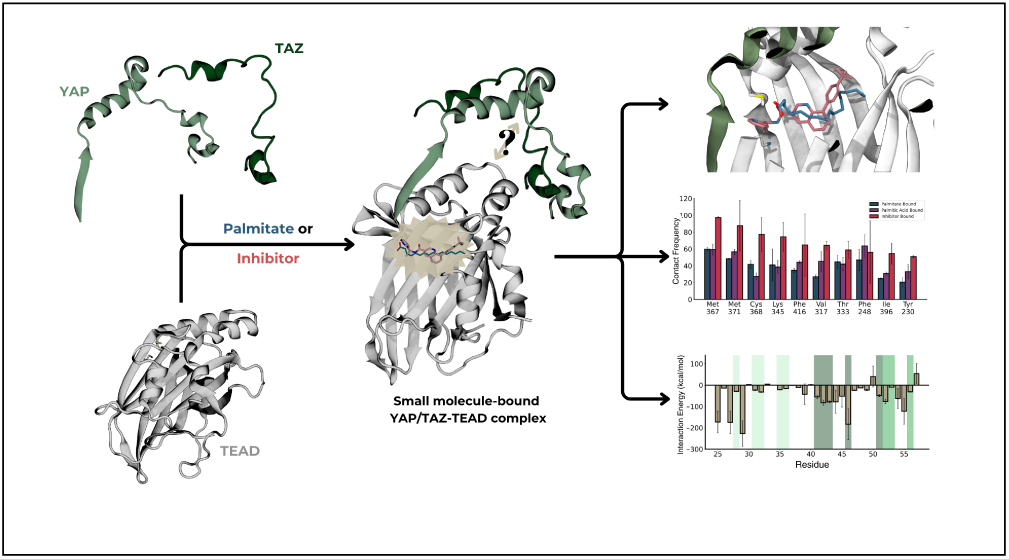

## References

(1) Misra, J. R.; Irvine, K. D. The Hippo Signaling Network and Its Biological Functions. Annual Review of Genetics 2018, 52, 65–87, PMID: 30183404.

(2) Cunningham, R.; Hansen, C. The Hippo pathway in cancer: YAP/TAZ and TEAD as therapeutic targets in cancer. Clinical Science 2022, 136, 197–222.

(3) Dong, J.; Feldmann, G.; Huang, J.; Wu, S.; Zhang, N.; Comerford, S. A.; Gayyed, M.; Anders, R. A.; Maitra, A.; Pan, D. Elucidation of a Universal Size-Control Mechanism in Drosophila and Mammals. Cell 2007, 130, 1120–1133.

(4) Basu, S.; Totty, N. F.; Irwin, M. S.; Sudol, M.; Downward, J. Akt Phosphorylates the Yes-Associated Protein, YAP, to Induce Interaction with 14-3-3 and Attenuation of p73-Mediated Apoptosis. Molecular Cell 2003, 11, 11–23.

(5) Lou, J.; Lu, Y.; Cheng, J.; Zhou, F.; Yan, Z.; Zhang, D.; Meng, X.; Zhao, Y. A chemical perspective on the modulation of TEAD transcriptional activities: Recent progress, challenges, and opportunities. European Journal of Medicinal Chemistry 2022, 243, 114684.

(6) Yu, F.-X.; Zhao, B.; Guan, K.-L. Hippo Pathway in Organ Size Control, Tissue Homeostasis, and Cancer. Cell 2015, 163, 811–828.

(7) Gridnev, A.; Maity, S.; Misra, J. R. Structure-based discovery of a novel small-molecule inhibitor of TEAD palmitoylation with anticancer activity. Frontiers in Oncology 2022, 12.

(8) Kaan, H. Y. K.; Chan, S. W.; Tan, S. K. J.; Guo, F.; Lim, C. J.; Hong, W.; Song, H. Crystal structure of TAZ-TEAD complex reveals a distinct interaction mode from that of YAP-TEAD complex. Scientific Reports 2017, 7, 2035.

(9) Pobbati, A.; Chan, S.; Lee, I.; Song, H.; Hong, W. Structural and Functional Similarity between the Vgll1-TEAD and the YAP-TEAD Complexes. Structure 2012, 20, 1135–1140.

(10) Li, Z.; Zhao, B.; Wang, P.; Chen, F.; Dong, Z.; Yang, H.; Guan, K.-L.; Xu, Y. Structural insights into the YAP and TEAD complex. Genes & Development 2010, 24, 235–240.

(11) Mesrouze, Y.; Bokhovchuk, F.; Meyerhofer, M.; Zimmermann, C.; Fontana, P.; Erdmann, D.; Chène, P. Study of the TEAD-binding domain of the YAP protein from animal species. Protein Science 2021, 30, 339–349.

(12) Noland, C.; Gierke, S.; Schnier, P.; Murray, J.; Sandoval, W.; Sagolla, M.; Dey, A.; Hannoush, R.; Fairbrother, W.; Cunningham, C. Palmitoylation of TEAD Transcription Factors Is Required for Their Stability and Function in Hippo Pathway Signaling. Structure 2016, 24, 179–186.

(13) Linder, M. E.; Deschenes, R. J. Palmitoylation: policing protein stability and traffic. Nature Reviews Molecular Cell Biology 2007, 8, 74–84.

(14) Chan, P.; Han, X.; Zheng, B.; DeRan, M.; Yu, J.; Jarugumilli, G. K.; Deng, H.; Pan, D.; Luo, X.; Wu, X. Autopalmitoylation of TEAD proteins regulates transcriptional output of the Hippo pathway. Nature Chemical Biology 2016, 12, 282–289.

(15) Tang, T. T.; Konradi, A. W.; Feng, Y.; Peng, X.; Ma, M.; Li, J.; Yu, F.-X.; Guan, K.- L.; Post, L. Small Molecule Inhibitors of TEAD Auto-palmitoylation Selectively Inhibit Proliferation and Tumor Growth of NF2-deficient Mesothelioma. Molecular Cancer Therapeutics 2021, 20, 986–998.

(16) Mesrouze, Y.; Meyerhofer, M.; Bokhovchuk, F.; Fontana, P.; Zimmermann, C.; Martin, T.; Delaunay, C.; Izaac, A.; Kallen, J.; Schmelzle, T.; others Effect of the acylation of TEAD4 on its interaction with co-activators YAP and TAZ. Protein science 2017, 26, 2399–2409.

(17) Li, Y.; Liu, S.; Ng, E. Y.; Li, R.; Poulsen, A.; Hill, J.; Pobbati, A. V.; Hung, A. W.; Hong, W.; Keller, T. H.; others Structural and ligand-binding analysis of the YAP-binding domain of transcription factor TEAD4. Biochemical Journal 2018, 475, 2043–2055.

(18) Mesrouze, Y.; Aguilar, G.; Meyerhofer, M.; Bokhovchuk, F.; Zimmermann, C.; Fontana, P.; Vissières, A.; Voshol, H.; Erdmann, D.; Affolter, M.; others The role of lysine palmitoylation/myristoylation in the function of the TEAD transcription factors. Scientific Reports 2022, 12, 4984.

(19) Noritsugu, K.; Suzuki, T.; Dodo, K.; Ohgane, K.; Ichikawa, Y.; Koike, K.; Morita, S.; Umehara, T.; Ogawa, K.; Sodeoka, M.; others Lysine long-chain fatty acylation regulates the TEAD transcription factor. Cell Reports 2023, 42.

(20) Liberelle, M.; Toulotte, F.; Renault, N.; Gelin, M.; Allemand, F.; Melnyk, P.; Guichou, J.-F.; Cotelle, P. Toward the Design of Ligands Selective for the C-Terminal Domain of TEADs. Journal of Medicinal Chemistry 2022, 65, 5926–5940, PMID: 35389210.

(21) Pashkovskaya, A. A.; Vazdar, M.; Zimmermann, L.; Jovanovic, O.; Pohl, P.; Pohl, E. E. Mechanism of long-chain free fatty acid protonation at the membrane-water interface. Biophysical Journal 2018, 114, 2142–2151.

(22) Waterhouse, A.; Bertoni, M.; Bienert, S.; Studer, G.; Tauriello, G.; Gumienny, R.; Heer, F. T.; de Beer, T. A.; Rempfer, C.; Bordoli, L.; Lepore, R.; Schwede, T. SWISS-MODEL: homology modelling of protein structures and complexes. Nucleic Acids Research 2018, 46, W296–W303.

(23) Anandakrishnan, R.; Aguilar, B.; Onufriev, A. V. H++ 3.0: automating pK prediction and the preparation of biomolecular structures for atomistic molecular modeling and simulations. Nucleic Acids Research 2012, 40, W537–W541.

(24) Myers, J.; Grothaus, G.; Narayanan, S.; Onufriev, A. A simple clustering algorithm can be accurate enough for use in calculations of pKs in macromolecules. Proteins: Structure, Function, and Bioinformatics 2006, 63, 928–938.

(25) Gordon, J. C.; Myers, J. B.; Folta, T.; Shoja, V.; Heath, L. S.; Onufriev, A. H++: a server for estimating p Ka s and adding missing hydrogens to macromolecules. Nucleic Acids Research 2005, 33, W368–W371.

(26) Case, D.; Belfon, K.; Ben-Shalom, I.; Brozell, S.; Cerutti, D.; Cheatham III, T.; Cruzeiro, V.; Darden, T.; Duke, R.; Giambasu, G.; Gilson, M.; Gohlke, H.; Goetz, A.; Harris, R.; Izadi, S.; Izmailov, S.; Kasavajhala, K.; Kovalenko, A.; Krasny, R.; Kurtzman, T.; Lee, T.; LeGrand, S.; Li, P.; Lin, C.; Liu, J.; Luchko, T.; Luo, R.; Man, V.; Merz, K.; Miao, Y.; Mikhailovskii, O.; Monard, G.; Nguyen, H.; Onufriev, A.; Pan, F.; Pantano, S.; Qi, R.; Roe, D.; Roitberg, A.; Sagui, C.; Schott-Verdugo, S.; Shen, J.; Simmerling, C.; Skrynnikov, N.; N.R.; Smith, J.; Swails, J.; Walker, R.; Wang, J.; Wilson, L.; Wolf, R.; Wu, X.; Xiong, Y.; Xue, Y.; York, D.; Kollman, P. AMBER 2020. 2020,

(27) Humphrey, W.; Dalke, A.; Schulten, K. VMD – Visual Molecular Dynamics. Journal of Molecular Graphics 1996, 14, 33–38.

(28) Tian, C.; Kasavajhala, K.; Belfon, K. A. A.; Raguette, L.; Huang, H.; Migues, A. N.; Bickel, J.; Wang, Y.; Pincay, J.; Wu, Q.; Simmerling, C. ff19SB: Amino-Acid-Specific Protein Backbone Parameters Trained against Quantum Mechanics Energy Surfaces in Solution. Journal of Chemical Theory and Computation 2020, 16, 528–552, PMID: 31714766.

(29) Izadi, S.; Anandakrishnan, R.; Onufriev, A. V. Building Water Models: A Different Approach. The Journal of Physical Chemistry Letters 2014, 5, 3863–3871, PMID: 25400877.

(30) Götz, A. W.; Williamson, M. J.; Xu, D.; Poole, D.; Le Grand, S.; Walker, R. C. Routine Microsecond Molecular Dynamics Simulations with AMBER on GPUs. 1. Generalized Born. Journal of Chemical Theory and Computation 2012, 8, 1542–1555, PMID: 22582031.

(31) Salomon-Ferrer, R.; Götz, A. W.; Poole, D.; Le Grand, S.; Walker, R. C. Routine Microsecond Molecular Dynamics Simulations with AMBER on GPUs. 2. Explicit Solvent Particle Mesh Ewald. Journal of Chemical Theory and Computation 2013, 9, 3878–3888, PMID: 26592383.

(32) Loncharich, R. J.; Brooks, B. R.; Pastor, R. W. Langevin dynamics of peptides: The frictional dependence of isomerization rates of N-acetylalanyl-N’-methylamide. Biopolymers 1992, 32, 523–535.

(33) Ryckaert, J.-P.; Ciccotti, G.; Berendsen, H. J. Numerical integration of the cartesian equations of motion of a system with constraints: molecular dynamics of n-alkanes. Journal of Computational Physics 1977, 23, 327–341.

(34) Berendsen, H. J. C.; Postma, J. P. M.; van Gunsteren, W. F.; DiNola, A.; Haak, J. R. Molecular dynamics with coupling to an external bath. The Journal of Chemical Physics 1984, 81, 3684–3690.

(35) Roe, D. R.; Cheatham, T. E. PTRAJ and CPPTRAJ: Software for Processing and Analysis of Molecular Dynamics Trajectory Data. Journal of Chemical Theory and Computation 2013, 9, 3084–3095, PMID: 26583988.

(36) Adasme, M. F.; Linnemann, K. L.; Bolz, S. N.; Kaiser, F.; Salentin, S.; Haupt, V.; Schroeder, M. PLIP 2021: expanding the scope of the protein–ligand interaction profiler to DNA and RNA. Nucleic Acids Research 2021, 49, W530–W534.

(37) Leddin, E.; Group, C. R.; Cisneros, G. A. CisnerosResearch/AMBER-EDA: First Release. 2021,

(38) Graham, S. E.; Syeda, F.; Cisneros, G. A. Computational Prediction of Residues Involved in Fidelity Checking for DNA Synthesis in DNA Polymerase I. Biochemistry 2012, 51, 2569–2578, PMID: 22397306.

(39) Torabifard, H.; Cisneros, G. A. Computational investigation of O2 diffusion through an intra-molecular tunnel in AlkB; influence of polarization on O2 transport. Chem. Sci. 2017, 8, 6230–6238.

(40) Torabifard, H.; Cisneros, G. A. Insight into wild-type and T1372E TET2-mediated 5hmC oxidation using ab initio QM/MM calculations. Chem. Sci. 2018, 9, 8433–8445.

(41) Maghsoud, Y.; Dong, C.; Cisneros, G. A. Computational Characterization of the Inhibition Mechanism of Xanthine Oxidoreductase by Topiroxostat. ACS Catalysis 2023, 13, 6023–6043.

(42) Chan, S. W.; Lim, C. J.; Loo, L. S.; Chong, Y. F.; Huang, C.; Hong, W. TEADs Mediate Nuclear Retention of TAZ to Promote Oncogenic Transformation*. Journal of Biological Chemistry 2009, 284, 14347–14358.

(43) Zhang, H.; Liu, C.-Y.; Zha, Z.-Y.; Zhao, B.; Yao, J.; Zhao, S.; Xiong, Y.; Lei, Q.-Y.; Guan, K.-L. TEAD transcription factors mediate the function of TAZ in cell growth and epithelial-mesenchymal transition. Journal of biological chemistry 2009, 284, 13355–13362.

(44) Pobbati, A. V.; Han, X.; Hung, A. W.; Weiguang, S.; Huda, N.; Chen, G.-Y.; Kang, C.; Chia, C. S. B.; Luo, X.; Hong, W.; others Targeting the central pocket in human transcription factor TEAD as a potential cancer therapeutic strategy. Structure 2015, 23, 2076–2086.

(45) Lu, W.; Wang, J.; Li, Y.; Tao, H.; Xiong, H.; Lian, F.; Gao, J.; Ma, H.; Lu, T.; Zhang, D.; others Discovery and biological evaluation of vinylsulfonamide derivatives as highly potent, covalent TEAD autopalmitoylation inhibitors. European Journal of Medicinal Chemistry 2019, 184, 111767.

(46) Bum-Erdene, K.; Zhou, D.; Gonzalez-Gutierrez, G.; Ghozayel, M. K.; Si, Y.; Xu, D.; Shannon, H. E.; Bailey, B. J.; Corson, T. W.; Pollok, K. E.; others Small-molecule covalent modification of conserved cysteine leads to allosteric inhibition of the TEAD Yap protein-protein interaction. Cell chemical biology 2019, 26, 378–389.

(47) Karatas, H.; Akbarzadeh, M.; Adihou, H.; Hahne, G.; Pobbati, A. V.; Yihui Ng, E.; Gueret, S. M.; Sievers, S.; Pahl, A.; Metz, M.; others Discovery of covalent inhibitors targeting the transcriptional enhanced associate domain central pocket. Journal of Medicinal Chemistry 2020, 63, 11972–11989.

(48) Holden, J. K.; Crawford, J. J.; Noland, C. L.; Schmidt, S.; Zbieg, J. R.; Lacap, J. A.; Zang, R.; Miller, G. M.; Zhang, Y.; Beroza, P.; others Small molecule dysregulation of TEAD lipidation induces a dominant-negative inhibition of hippo pathway signaling. Cell reports 2020, 31.

(49) Callus, B. A.; Finch-Edmondson, M. L.; Fletcher, S.; Wilton, S. D. YAPping about and not forgetting TAZ. FEBS Letters 2019, 593, 253–276.

(50) Wang, H.; Wang, J.; Zhang, S.; Jia, J.; Liu, X.; Zhang, J.; Wang, P.; Song, X.; Che, L.; Liu, K.; Ribback, S.; Cigliano, A.; Evert, M.; Wu, H.; Calvisi, D. F.; Zeng, Y.; Chen, X. Distinct and Overlapping Roles of Hippo Effectors YAP and TAZ During Human and Mouse Hepatocarcinogenesis. Cellular and Molecular Gastroenterology and Hepatology 2021, 11, 1095–1117.

(51) Fallah, S.; Beaulieu, J.-F. Differential influence of YAP1 and TAZ on differentiation of intestinal epithelial cell: A review. The Anatomical Record 2023, 306, 1054–1061.

(52) LeBlanc, L.; Ramirez, N.; Kim, J. Context-dependent roles of YAP/TAZ in stem cell fates and cancer. Cellular and Molecular Life Sciences 2021, 78, 4201–4219.

(53) Reggiani, F.; Gobbi, G.; Ciarrocchi, A.; Sancisi, V. YAP and TAZ Are Not Identical Twins. Trends in Biochemical Sciences 2021, 46, 154–168.

