## Supplementary Tables 1-2, Supplementary Figures 1-6 for "Allosteric Modulation of YAP/TAZ-TEAD Interaction by Palmitoylation and Small Molecule Inhibitors"

| Residue | PLT-apo | PLM-apo | INH-apo |
|---|---|---|---|
| **52** | -1.6788 | -1.2576 | 0.0815 |
| **53** | -1.4074 | 3.7443 | -17.8970 |
| **54** | -14.2839 | -9.1088 | -9.1371 |
| **55** | -4.9157 | -1.8812 | -3.7765 |
| **56** | -12.0898 | 4.6284 | 7.5205 |
| **57** | 2.4522 | 0.5148 | -1.6458 |
| **58** | -20.6638 | -67.9862 | -0.2532 |
| **59** | -4.1113 | -6.0169 | -3.3921 |
| **60** | 54.2961 | 40.4970 | 3.6644 |
| **61** | 8.4537 | -4.0026 | -13.8957 |
| **62** | 51.9963 | 33.1884 | 26.2896 |
| **63** | 0.1333 | -1.0259 | 0.6105 |
| **64** | 53.7006 | 58.5977 | 15.4675 |
| **65** | 14.6919 | 12.0511 | 9.4502 |
| **66** | 52.1783 | 29.8723 | -14.7268 |
| **67** | 0.5366 | 1.8074 | 0.5065 |
| **68** | 14.1376 | 13.9232 | 2.2608 |
| **69** | 7.5830 | 6.0963 | 1.7268 |
| **70** | -1.7232 | 1.6037 | 0.5068 |
| **71** | -0.6217 | 1.9859 | 2.5615 |
| **72** | -2.7969 | -4.4482 | -4.0036 |
| **73** | -14.6185 | -5.8849 | -2.3780 |
| **74** | -0.4861 | -0.5159 | 0.4059 |
| **75** | 2.0304 | 1.9732 | 1.8833 |
| **76** | -27.5154 | -36.5959 | -32.1182 |
| **77** | 1.6235 | 1.6138 | 1.2560 |
| **78** | 3.6211 | 0.9556 | 3.6467 |
| **79** | -2.5854 | -4.0981 | 1.2114 |
| **80** | 2.5210 | 0.8706 | 4.0196 |
| **81** | 0.3668 | 0.9656 | -3.2424 |
| **82** | -1.0947 | 1.9931 | -5.9805 |
| **83** | 5.8005 | 10.2715 | 3.3907 |
| **84** | 1.0217 | 1.0708 | -5.8746 |
| **85** | 13.7281 | 10.0497 | -9.0677 |
| **86** | 15.9693 | 13.5716 | -12.7852 |
| **87** | 18.6775 | -12.6946 | -103.6946 |
| **88** | 3.6280 | 1.2609 | 0.4835 |
| **89** | 51.4690 | 28.5870 | -16.5120 |
| **90** | -26.9683 | -28.9203 | -27.7043 |
| **91** | -1.4835 | -0.3716 | 0.2993 |
| **92** | 9.3529 | 2.5953 | 2.8775 |
| **93** | 29.8762 | 27.7606 | 27.4264 |
| **94** | 15.1249 | 1.1099 | 5.3394 |
| **95** | 5.4559 | 1.6656 | -0.8712 |
| **96** | -0.5693 | 1.7128 | -0.2025 |
| **97** | -38.8528 | -32.7147 | -32.3816 |
| **98** | 10.8180 | 0.3681 | 2.3305 |
| **99** | 10.7687 | 3.9025 | 3.7454 |
| **100** | 31.7272 | 31.2003 | 39.6227 |
| **Total** | **315.2738** | **134.4856** | **-152.9559** |

Table S1: EDA analysis between individual YAP residues and the combined TEAD protein. For each residue, the delta energy is shown between the systems with palmitate (PLT), palmitic acid (PLM), or the inhibitor (INH) bound and the apo system to show if the binding of the small molecule is stabilizing or destabilizing the interaction between YAP and TEAD. Values are based on averages from all three trials of each, over the entirety of the simulation, and are displayed as mean ($\pm$ standard deviation).

| **TAZ Residue** | **PLT-apo** | **PLM-apo** | **INH-apo** |
|---|---|---|---|
| **25** | 41.9723 | 70.8846 | 108.8100 |
| **26** | -0.9073 | -1.7501 | 5.4646 |
| **27** | 39.3880 | 125.3457 | 127.2684 |
| **28** | -0.0437 | 7.4133 | 13.8174 |
| **29** | 47.3702 | 120.0313 | 170.7043 |
| **30** | 0.9827 | -1.2011 | -13.4356 |
| **31** | 9.8784 | 13.0517 | -8.3315 |
| **32** | 0.6662 | 9.6360 | -10.1612 |
| **33** | 0.8285 | -3.4234 | -11.0899 |
| **34** | -4.7200 | 0.7237 | -0.9881 |
| **35** | 4.6295 | 14.3827 | 18.2220 |
| **36** | -6.7934 | 5.6418 | 9.3763 |
| **37** | -5.5540 | -2.1709 | -4.9399 |
| **38** | 0.3335 | -0.5987 | -2.7434 |
| **39** | -5.1159 | 35.0862 | -21.6404 |
| **40** | -7.2121 | -6.1647 | -14.7209 |
| **41** | 37.2530 | 29.0127 | 25.4966 |
| **42** | 79.5157 | 69.2866 | 51.8518 |
| **43** | 12.5044 | 30.0285 | 57.2399 |
| **44** | -127.1043 | -44.9407 | -33.4168 |
| **45** | -22.1078 | 7.8644 | 9.0000 |
| **46** | -15.3364 | 91.9666 | 102.6472 |
| **47** | 9.7256 | 10.6770 | 17.3607 |
| **48** | 0.9218 | 1.0871 | 3.7012 |
| **49** | -3.7550 | 10.7033 | 14.0169 |
| **50** | 37.2956 | 2.8923 | -5.8454 |
| **51** | 0.2385 | 21.4851 | 27.7641 |
| **52** | -1.2533 | 20.9693 | 51.6797 |
| **53** | -1.8029 | 3.8404 | 7.3934 |
| **54** | -38.5781 | -26.2763 | 7.1694 |
| **55** | 146.0850 | 167.0517 | 167.8172 |
| **56** | -4.4099 | 17.2561 | 24.9466 |
| **57** | 33.2376 | -20.5615 | -4.9673 |
| **Total** | **258.1325** | **779.2309** | **889.4672** |

Table S2: EDA analysis between individual TAZ residues and the combined TEAD protein. For each residue, the delta energy is shown between the systems with palmitate (PLT), palmitic acid (PLM), or the inhibitor (INH) bound and the apo system to show if the binding of the small molecule is stabilizing or destabilizing the interaction between TAZ and TEAD. Values are based on averages from all three trials of each, over the entirety of the simulation, and are displayed as mean ($\pm$ standard deviation). Palmitic acid-bound trial that showed dissociation is omitted as EDA analysis gave infinite values.

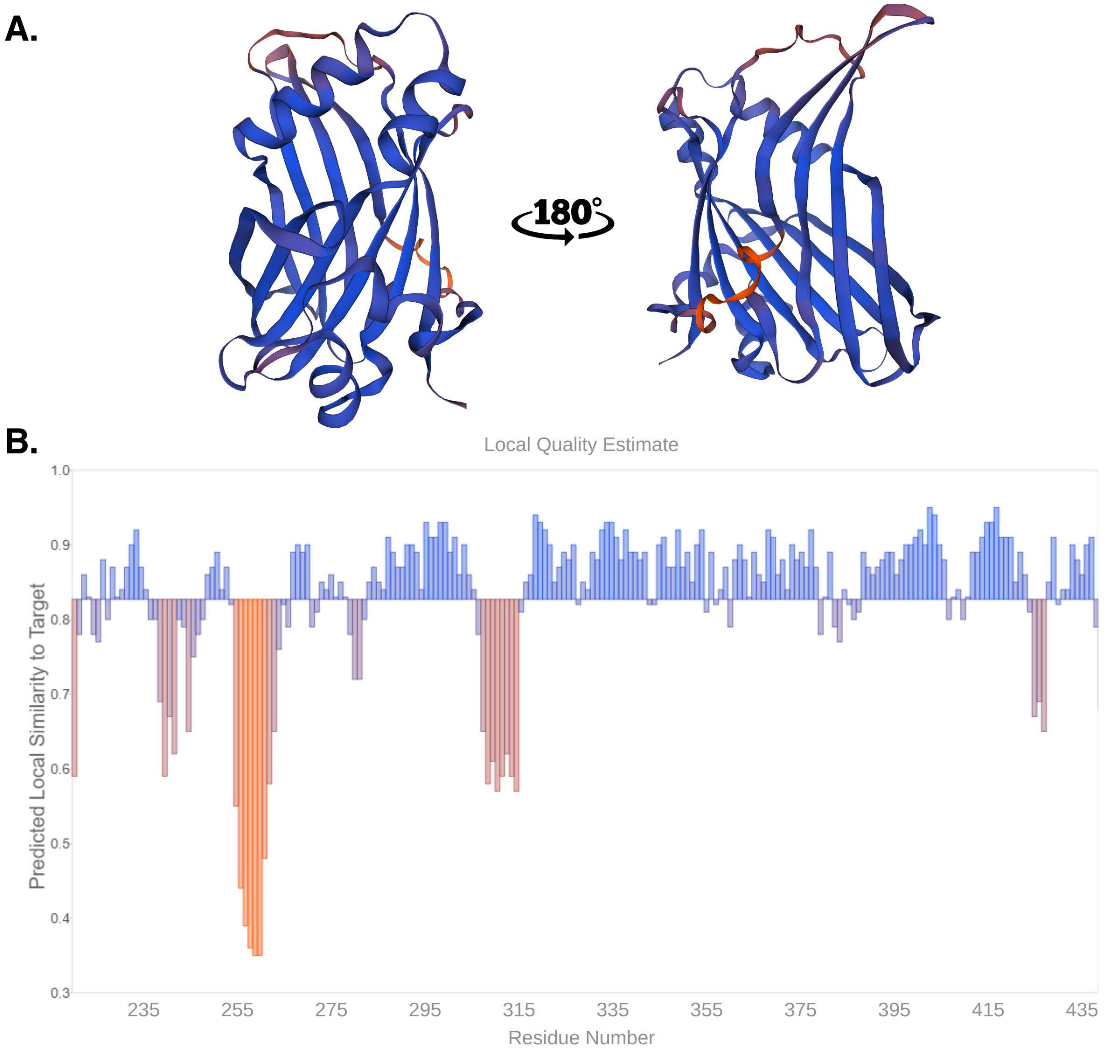


Figure S1: Homology modeling results for TEAD Yap-Binding Domain used to add residues 253-260 which were missing from the crystal structure used (PDB ID: 7CNL). The per-residue model quality score as determined by SWISS-MODEL are A. shown on protein snapshots and B. plotted with each residue of the model (x-axis) and the expected similarity to the native structure (y-axis).

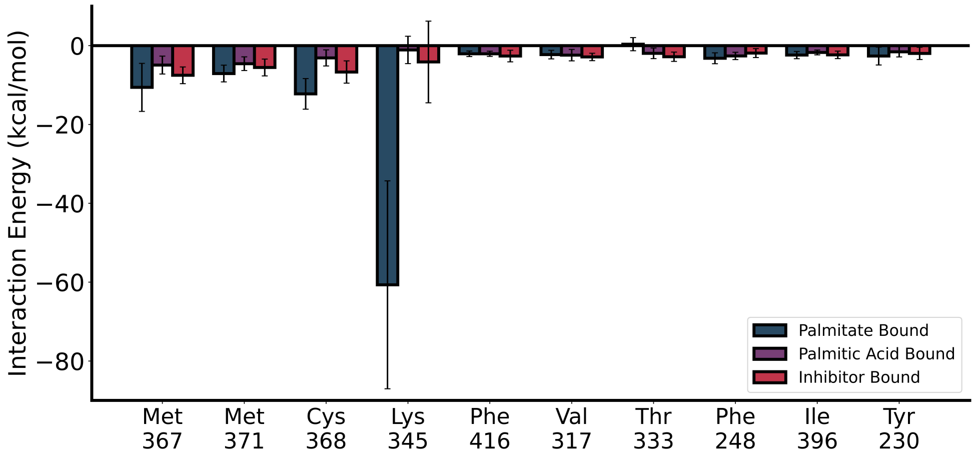


Figure S2: Average interaction energy between each of the small molecules and the residues lining the palmitate binding pocket. Error bars show the standard deviation between the three trials.

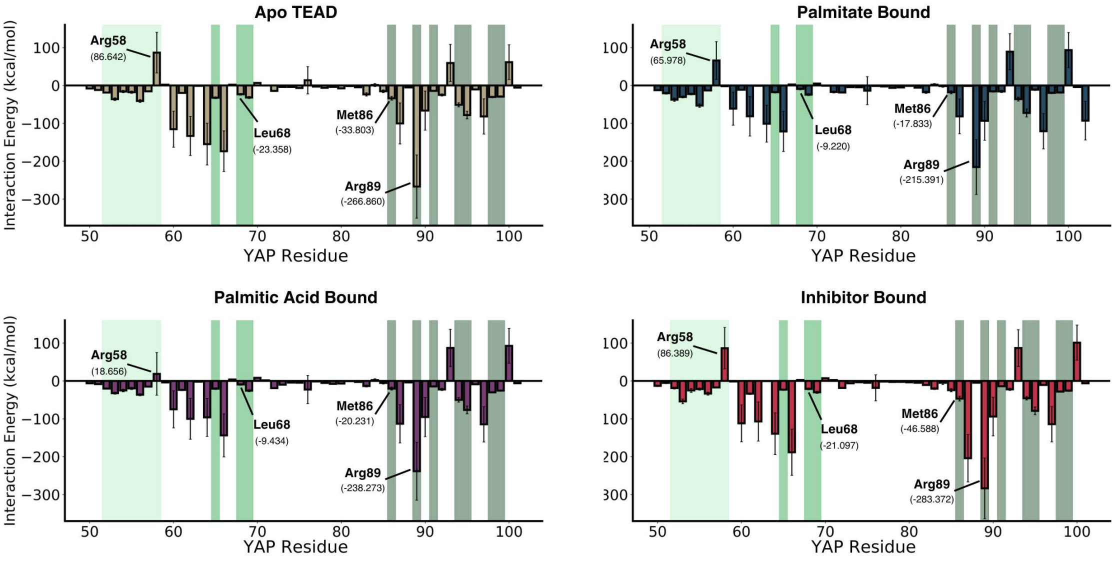


Figure S3: Energy Decomposition Analysis (EDA) showing the combined interaction energy (van der Waals and electrostatics) for each YAP residue with all of the TEAD residues. Error bars show the standard deviation between the three trials. Distinct shaded regions highlight the three binding interfaces. The lightest shade of green marks Interface 1 (residues 52-58), an intermediate shade of green indicates Interface 2 (residues 65, 68, 69), and the darkest shade represents Interface 3 (residues 86, 89, 91, 94-95, 98-99).

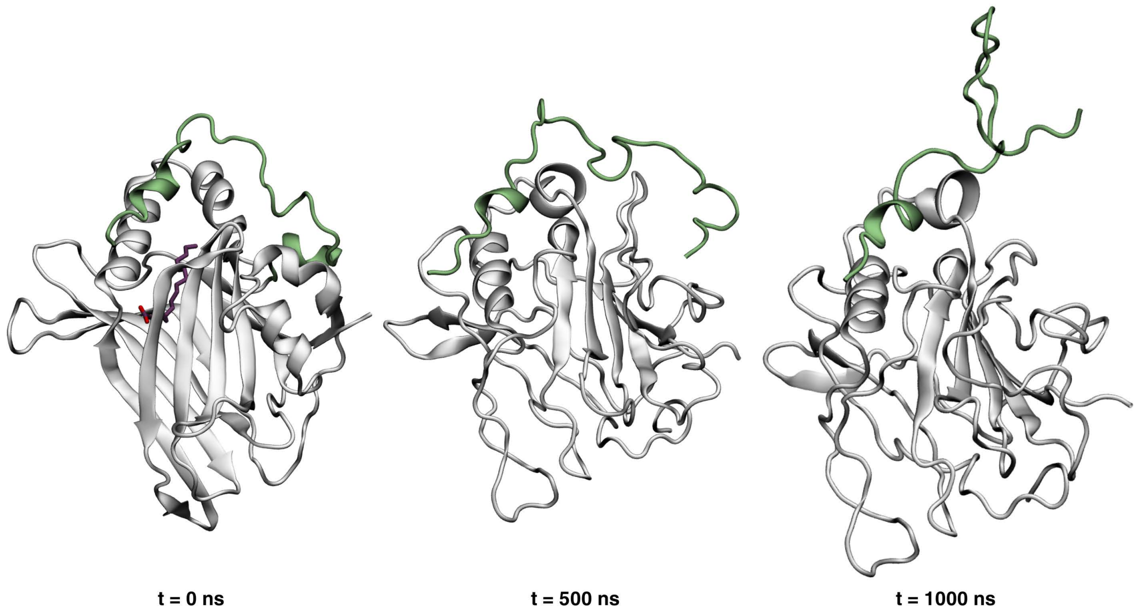


Figure S4: Simulations snapshots of the omitted palmitic acid-bound TAZ-TEAD heterodimer trial, showing the palmitic acid leaving the PBP and TAZ beginning to dissociate from TEAD. TEAD protein is shown in white, TAZ protein is shown in green, and palmitic acid is shown in purple.

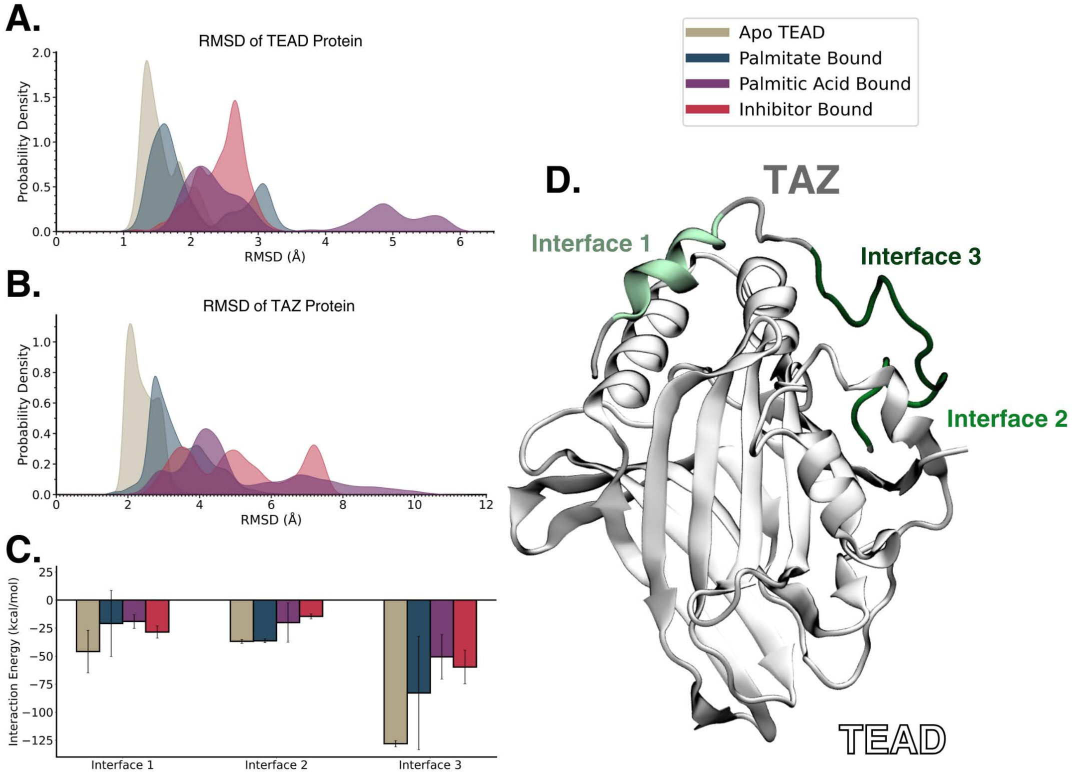


Figure S5: Combined results for the TAZ-TEAD heterodimer simulations, including palmitic acid-bound trial which showed palmitic acid leaving the PBP and TAZ becoming dissociated from TEAD. A. RMSD of the backbone atoms of TEAD protein in each of the simulations. B. RMSD of the backbone atoms of TAZ protein in each of the simulations. C. Total interaction energy between the residues comprising each of the binding interfaces of TAZ and TEAD. Error bars show the standard deviation between the three trials. D. Visualization of the three binding interfaces between TAZ and TEAD.

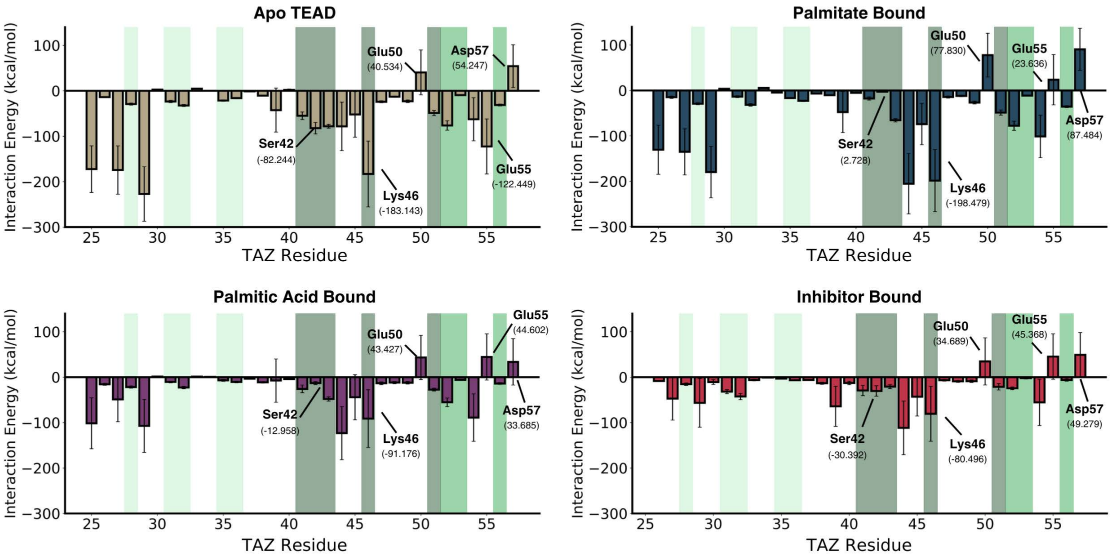


Figure S6: Energy Decomposition Analysis (EDA) showing the combined interaction energy (van der Waals and electrostatics) for each TAZ residue with all of the TEAD residues. Error bars show the standard deviation between the three trials. Distinct shaded regions highlight the three binding interfaces. The lightest shade of green marks Interface 1 (residues 28, 31-32, 35-36), an intermediate shade of green indicates Interface 2 (residues 52, 53, 56), and the darkest shade represents Interface 3 (residues 41-43, 46, 51). Palmitic acid-bound trial that showed dissociation is omitted as EDA analysis gave infinite values.
